# The genome of the endangered *Macadamia jansenii* displays little diversity but represents an important genetic resource for plant breeding

**DOI:** 10.1101/2021.09.08.459545

**Authors:** Priyanka Sharma, Valentine Murigneux, Jasmine Haimovitz, Catherine J. Nock, Wei Tian, Ardashir Kharabian Masouleh, Bruce Topp, Mobashwer Alam, Agnelo Furtado, Robert J. Henry

## Abstract

Macadamia, a recently domesticated expanding nut crop in the tropical and subtropical regions of the world, is one of the most economically important genera in the diverse and widely adapted Proteaceae family. All four species of *Macadamia* are rare in the wild with the most recently discovered, *M. jansenii*, being endangered. The *M. jansenii* genome has been used as a model for testing sequencing methods using a wide range of long read sequencing techniques. Here we report a chromosome level genome assembly, generated using a combination of Pacific Biosciences sequencing and Hi-C, comprising 14 pseudo-molecules, with a N50 of 58 Mb and a total 758 Mb genome assembly size of which 56% is repetitive. Completeness assessment revealed that the assembly covered 96.9% of the conserved single copy genes. Annotation predicted 31,591 protein coding genes and allowed the characterization of genes encoding biosynthesis of cyanogenic glycosides, fatty acid metabolism and anti-microbial proteins. Re-sequencing of seven other genotypes confirmed low diversity and low heterozygosity within this endangered species. Important morphological characteristics of this species such as small tree size and high kernel recovery suggest that *M. jansenii* is an important source of these commercial traits for breeding. As a member of a small group of families that are sister to the core eudicots, this high-quality genome also provides a key resource for evolutionary and comparative genomics studies.

## Introduction

Macadamia is a recent domesticate with a complex domestication history (Peace, 2005). The four currently recognised *Macadamia* species are endemic to the central coast of eastern Australia (Mast et al., 2008). However, macadamia was first domesticated in Hawaii around 100 years ago, with most of the global production based upon the Hawaiian domesticated germplasm (Hardner, 2016). Macadamia is a member of the Proteaceae family, one of a group of families that are a sister to the core eudicots (Gross and Weston, 1992; Christenhusz and Byng, 2016). Macadamia is the first Australian native plant that has been widely grown as a food plant (Peace et al., 2013). All of the Hawaiian macadamia cultivars has been reported to be based upon only a few or possibly even a single tree from Australia (Nock et al., 2019). This resulting narrow gene pool makes it susceptible to disease and climate change, whereas the unexploited wild macadamia germplasm of Australia provides an opportunity for great improvement of this newly domesticated crop. Despite a rapid international increase in macadamia production, breeding is restricted because of lack of genomic information (Topp et al., 2019).

Macadamia is the most widely grown Australian native food crop (Peace et al., 2013). Macadamia production was valued at USD 1.17 billion in 2019 and production is expected to grow at a rate of 9.2% from 2020 to 2027 (https://www.grandviewresearch.com/industry-analysis/macadamia-nut-market). Among the macadamia species, *M. integrifolia*, the species from which most of the domesticated gene pool is derived (Hardner, 2016), was the first genome to be sequenced (Nock et al., 2016). This genome, of cultivar HAES 741, has supported initial efforts at genome based breeding (O’Connor et al., 2018) and has recently been upgraded to chromosome level with a contig N50 of 413 Kb. The other species that has been a contributor to domesticated germplasm, *M. tetraphylla*, has been sequenced with an N50 of 1.18 Mb (Niu et al., 2020).

All species are rare in the wild but *M. jansenii* is endangered and is only found in a limited area to the north-west of Bundaberg, Queensland (Shapcott and Powell, 2011; Hayward et al., 2021). *Macadamia jansenii* is endangered under the Australian (EPBC) Act and critically endangered under the Queensland (Qld Nature Conservation Act) legislation (Gross and Weston, 1992). Due to the expected low heterozygosity associated with the extremely small population size, this species has been used as a model to compare available genome sequencing technologies (Murigneux et al., 2020; Sharma et al., 2021). *Macadamia jansenii* has been sequenced (Murigneux et al., 2020), using three long read sequencing technologies, Oxford Nanopore (PromethION), PacBio (Sequel I) and BGI (Single-tube Long Fragment Read). The genome was recently updated by sequencing using the PacBio HiFi sequencing (Sharma et al., 2021). Here, we report chromosome level assembly of the same genotype using Hi-C and annotation of the genome. This provides a platform that allows analysis of key genes of importance in macadamia breeding, a reference genome in this group of angiosperms and insights into the impact of rarity on plant genomes.

This high quality reference genome also provides a platform for analysis of three unique attributes of macadamia, the high levels of unusual fatty acids (Hu et al., 2019b), high cyanogenic glucoside content, (Nock et al., 2016) and the presence of a novel anti-microbial peptide (Marcus et al., 1999). The fatty acid, palmitoleic acid (16:1) is found in large amounts in macadamia and has been considered to have potential human health benefits (Solà Marsiñach and Cuenca, 2019; Song et al., 2018). Cyanogenic glycosides in plants are part of their defence against herbivores. However, the highly bitter nuts of *M. jansenii* are not edible and use of this species in macadamia breeding will require selection to ensure high levels of cyanogenic glycosides are avoided. Identification of the associated genes could assist by providing molecular tools for use in breeding selection. A novel antimicrobial protein was reported in the kernals of *M. integrifolia* (Marcus et al., 1999). These small antimicrobial proteins were found to be produced by processing of a larger pre-cursor protein. As fungal infection and insect herbivores are major hurdles in macadamia production (Dahler et al., 1995; Nock et al., 2016; Marcus et al., 1999), retention of the antimicrobial protein and cyanogenesis in some parts of the plant may be important. Analysis of candidate genes for these traits may assist in understanding and manipulating in macadamia breeding.

## Results

### Genome sequencing and assembly

A pseudo-molecule level genome assembly of Pac Bio contigs (Murigneux et al., 2020) was produced using Hi-C. The estimated genome size of *M. jansenii* was 780 Mb (Murigneux et al., 2020) and the size of the final Hi-C assembly is 758 Mb comprised of 219 scaffolds with an N50 of 52Mb **(Table 1)**. Of this 97% was anchored to the 14 largest scaffolds representing the 14 chromosomes **(Figure S1, Table S1)**. Comparison of the PacBio assembly with the Hi-C chromosome assembly shows the number of scaffolds decreased from 762 to 219 and the length of the longest scaffold increased 6-fold **(Table 1)**. The L50 reduced from 135 to 7 scaffolds and the N50 was improved from 1.58 Mb to 52 Mb.

**Table 1.** *Macadamia jansenii* genome sequencing and assembly statistics.

|  | **PacBio** | **Dovetail Chicago** | **Dovetail Hi-C**<br>**assembly** |
| --- | --- | --- | --- |
| Library Statistics | 3,170,206 reads | 213M read pairs;<br>2x150 bp | 156M read pairs; 2x 150<br>bp |
| Coverage | 84 X | 88 X | 3,601 X |
| **Genome assembly** |  |  |  |
| Total Length | 758.28 Mb | 758.30 Mb | 758.43 Mb |
| L50/N50* | 135 scaffolds; 1.58<br>Mb | 199 scaffolds; 1.0<br>Mb | 7 scaffolds; 52.1Mb |
| L90/N90* | 457 scaffolds; 0.51<br>Mb | 767 scaffolds; 0.23<br>Mb | 13 scaffolds; 45.61 Mb |
| Longest Scaffold | 10,537,631 bp | 8,434,305 bp | 67,682,215 bp |
| Number of<br>Scaffolds | 762 | 1,529 | 219 |
| **BUSCO results\*** |  |  |  |
| Single genes | 79.10% | 80.10% | 80.80% |
| Duplicated genes | 17.60% | 17.10% | 16.30% |
| Fragmented genes | 0.90% | 1.00% | 1.00% |
| Missing genes | 2.00% | 2.00% | 2.10% |

\* Eudicots\_odb10 dataset, Number of BUSCOs= 2326.

### Assembly completeness and repeat element analysis

The completeness of the *M. jansenii* assembly was assessed by Benchmarking Universal Single-Copy orthologs (BUSCO) (Simão et al., 2015). This analysis revealed 96.9% complete genes (single and duplicated) in the Hi-C assembly **(Table 1)**. A total of 423.6 Mb, representing 55.9% of the Hi-C assembly was identified as repetitive **(Table 2)**. Class I TE (Transposable Elements) repeats were the most abundant repetitive elements representing 30% of the genome, including LTRs (24%), LINE (5.67%) and SINE (0%) and Class II TE repeats were 1.56%.

**Table 2.** Annotation of repeat sequences in the *M. jansenii* genome.

|  |  | Hi-C Assembly |
| --- | --- | --- |
| Total Repetitive content |  | 55.9% |
| Class I TEs repeats |  | 29.9% |
|  | LTRs | 24% |
|  | LINE | 5.67% |
|  | SINE | 0% |
| Class II TEs repeats |  | 1.56% |
| Low complexity repeats |  | 0.33% |
| Simple repeats |  | 1.35% |

### Structural and functional annotation

A total of 31,591 genes were identified in the repeat-masked Hi-C *M. jansenii* genome using an homology-based and RNA assisted approach. The average length of the genes was 1,368 bp **(Table 3)**. Of a total of 31,591 transcripts, only 22,500 sequences (71%) were annotated by BLAST2GO **(Figure S2)**. The transcripts were functionally annotated using Gene Ontology (GO) terms to assess the potential role of the genes in the *M. jansenii* genome. The most abundant *M. jansenii* specific gene families were organic cyclic and heterocyclic compound among the molecular function; organic and cellular metabolic among the biological process; and protein-containing binding membrane and intracellular organelle among the cellular component (**Figure S3**). The comparison of the three *Macadamia* genomes, assembled so far, showed *M. jansenii* has the highly continuous assembly with highest number of BUSCO genes **(Table 4)**.

**Table 3.** Genes predicted in the *M. jansenii* genome

| Gene prediction |  |
| --- | --- |
| Total number of genes | 31,591 |
| Total coding region | 43,235,907 bp |
| Average length of genes | 1,368 bp |
| Number of single-exon genes | 2,458 |
| Number of genes with annotation | 22,500 |
| Cyanogenic genes | 82 |
| Fatty acid genes | 47 |
| Anti-microbial genes | 1 |

**Table 4.** Comparison of genome assemblies of three *Macadamia* species.

|  | *M. integrifolia* (V1) | *M. integrifolia* (V2) | *M. tetraphylla* | *M. jansanii* |
| --- | --- | --- | --- | --- |
| Assembly length (Mb) | 518.49 | 744.64 | 750.53 | 758.43 |
| N50 (kb) | 4.7 | 413.4 | 1.2 | 52.1 |
| No. of contigs/scaffolds | 193,493 | 4094 | 4,335 | 219 |
| Repeats | 37.00% | 55.00% | 61.42% | 55.90% |
| BUSCO | 77.40% | 90.20% | 89.72% | 96.90% |
| No. of coding genes | 35,337 | 34,274 | 31,571 | 31,591 |

### Anti-microbial genes

Antimicrobial proteins have been reported in *M. integrifolia* (Marcus et al., 1999). In addition to antimicrobial properties these seed storage proteins are homologous to vicilin 7S globulins and have been identified as putative allergens (Rost et al., 2020; Rost et al., 2016). A cDNA sequence, from *M. integrifolia*, encoding these proteins, MiAMP-2, has been reported to contain four repeat segments, with each segment comprised of cysteine rich motifs (C-X-X-X-C-(10 to12) X-C-X-X-X-C), where X is any other amino acid residue (Marcus et al., 1999). Blast analysis identified homologues in the *M. jansenii* genome (**Figure S4**). The ANN01396 transcript from *M jansenii*, also showed four repeat segments of cysteine motifs with the same structure as found in MiAMP-2 **(Figure 1A)**. Comparison of the translated protein sequences indicated a high level of homology with only 28 differences in the 665 aa sequence **(Figure 1B)**. The *M. jansenii* sequence provides the first genomic sequence for this novel anti-microbial gene and reveals the presence of an intron in the 5’ UTR (**Figure S5)**.

**Figure 1.**
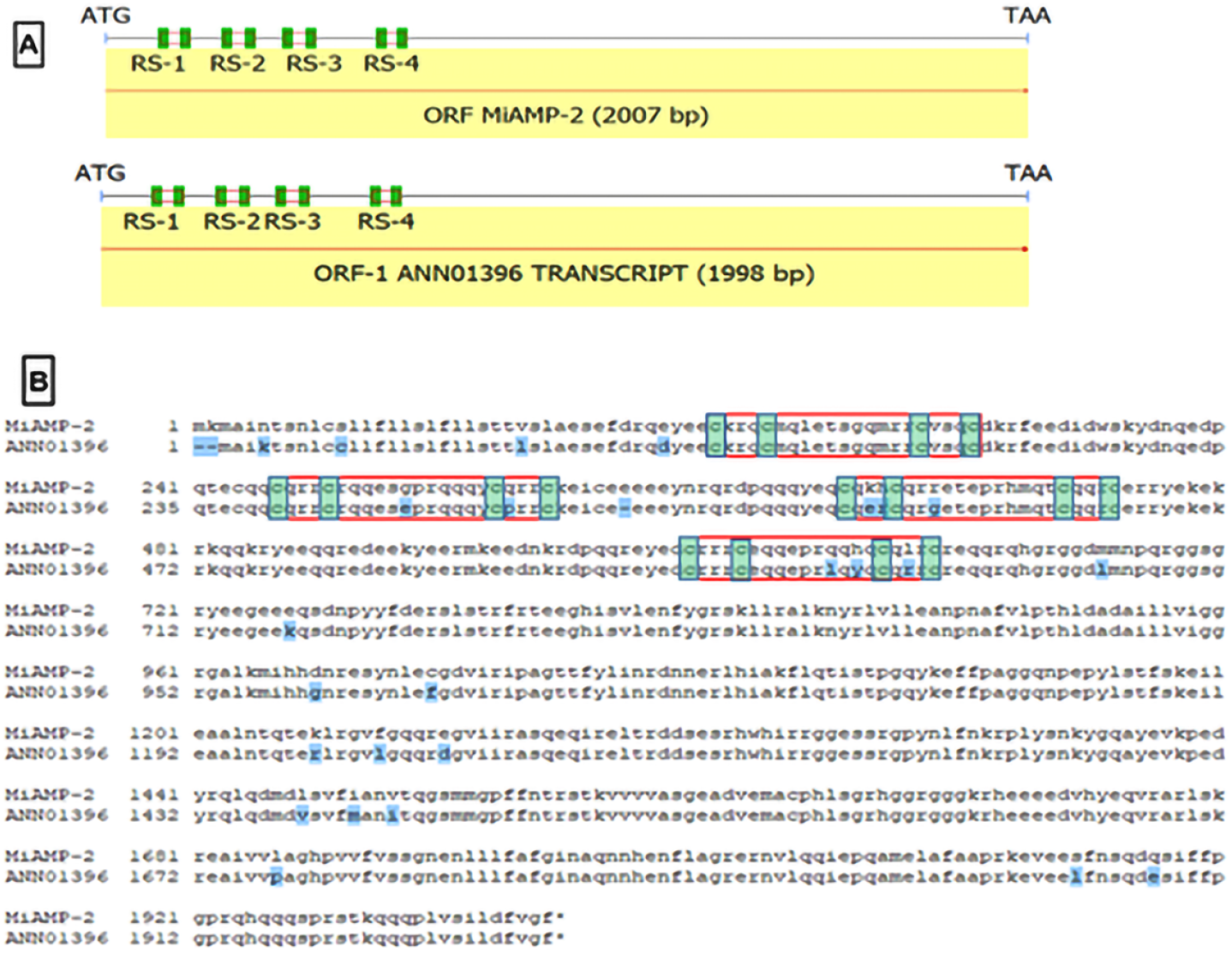
Anti-microbial peptide structure **Figure 1(A)** is the cDNA sequence of anti-microbial gene of *M. integrifolia* with four repeat segments (RS), shown in red open boxes and cysteine residues in green filled boxes aligned with *M. jansenii* transcript sequence ANN01396, showing same pattern. Figure 1(B) shows the alignment of the anti-microbial peptide sequence from the *M. integrifolia* and *M. jansenii*. The first half of the sequence shows the repeat segments within red boxes with green highlighted cysteine residues. Differences in amino acid sequence throughout the alignment as shown in blue highlighted text.

### Cyanogenic glycoside genes

*M. jansenii* has bitter nuts, presumably because of the presence of cyanogenic glycosides (Nock et al., 2016; Castada et al., 2020). Analysis of genes of cyanogenic glycoside metabolism detected a total of 76 putative genes in the *M. jansenii* genome. These genes were distributed throughout the genome (**Figure 2(A)**). The largest number of these genes (22) are encoded by UGT85 which is responsible for conversion of Hydronitrile to cyanogenic glucoside. In contrast only 14 genes for Cyp 79, the first gene in the pathway, was found (**Figure 2(B) & Table S8**).

**Figure 2.**
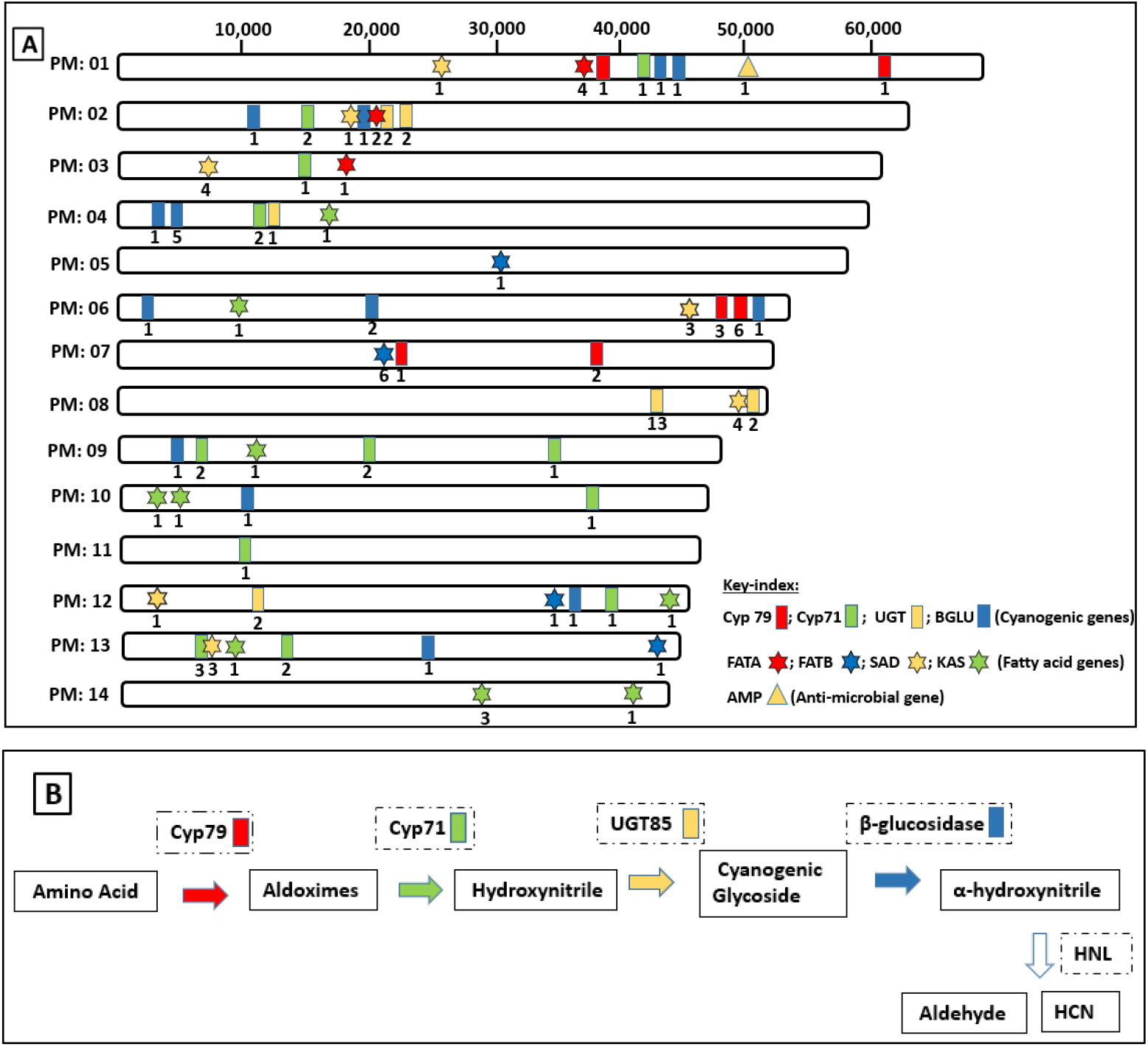
Pseudo-chromosomes of *M. jansenii* with location of cyanogenic, fatty acid and anti-microbial genes. **Figure 2(A)** putative cyanogenic, fatty acid and anti-microbial gene locations are shown on 14 pseudo molecules of *M. jansenii*. The bars show the cyanogenic genes, the stars show the genes involved in fatty acid pathway and the triangle shows the antimicrobial gene location on the pseudo-chromosome, the color key-index is given along with the figure. Pseudo-chromosomes are not to scale. **Figure 2(B)** illustrates the cyanogenic pathway and the main enzymes involved.

### Fatty acid metabolism genes

This study identified the key enzymes involved in fatty acid biosynthesis: elongases (e.g., KAS, FATA, FATB) and desaturases (e.g., SAD). A total of 44 of these genes were found in the *M. jansenii* genome. Stearoyl-ACP desaturases (SAD) which convert 18:0 to 18:1 was found to be abundant with 17 genes present (**Figure 2(A) & Table S7**).

### Heterozygosity and genetic diversity

To study the genetic diversity within the species, re-sequencing of seven other individuals was performed. A total of 166 M to 167 M reads of 150 bp in length were obtained. This represents a coverage of around 32 X of the *M. jansenii* genome. The seven accessions analysed had between 5.4 and 7.0 million variants relative to the reference genome (Table 5). Most of these were SNPs with less than 600,000 indels in all genotypes. Most SNPs were heterozygous with approximately 1 million or less homozygous SNP variants in each individual. The level of SNP heterozygosity for the 8 genotypes (including the reference) was found to be in the range of 0.26% to 0.34% with an average of 0.31 % (Table 5). The genotypes varied in their divergence from the reference with most unique variants being heterozygous and only 85,000 to 165,00 unique homozygous SNPs being found in an individual and not present in the other seven genotypes.

**Table 5.** Heterozygosity and genetic variation in *M. jansenii*

| Accession ID | Number<br>of<br>polymorp<br>hic sites | Number<br>of Indels | Number<br>of SNP | Variant<sup>1</sup><br>Positions:<br>Homozyg<br>ous SNPs | Variant<sup>1</sup><br>Positions:<br>Heterozygous<br>SNPs | SNP<br>Heterozy<br>gosity | Unique<sup>2</sup><br>Heterozygo<br>us variants | Unique<sup>2</sup><br>Homozygous<br>variants | Total<br>unique<sup>2</sup><br>polymorphi<br>c positions |
| --- | --- | --- | --- | --- | --- | --- | --- | --- | --- |
| 1005* | 5,418,086 | 486,846 | 4,764,835 | 4,019 | 2,428,956 | 0.31 | 2,249,732 | 3,070 | 2,252,802 |
| 1161004 | 5,415,612 | 377,580 | 4,901,611 | 784,323 | 2,038,553 | 0.26 | 1,902,705 | 111,100 | 2,013,805 |
| 1161003 | 6,785,189 | 555,641 | 6,034,679 | 1,047,938 | 2,465,089 | 0.32 | 2,306,771 | 162,541 | 2,469,312 |
| 1161005 | 6,204,994 | 531,550 | 5,488,354 | 780,593 | 2,347,362 | 0.30 | 2,190,169 | 85,258 | 2,275,427 |
| 1161001a | 6,977,842 | 574,625 | 6,196,254 | 875,565 | 2,649,035 | 0.34 | 2,484,209 | 109,875 | 2,594,084 |
| 1003 | 7,050,861 | 586,001 | 6,254,227 | 891,669 | 2,672,103 | 0.34 | 2,505,726 | 113,837 | 2,619,563 |
| 1002 | 6,759,260 | 586,334 | 5,973,425 | 1,044,208 | 2,447,418 | 0.31 | 2,286,675 | 165,048 | 2,451,723 |
| 1161001b | 6,704,384 | 548,292 | 5,962,434 | 824,632 | 2,556,695 | 0.33 | 2,394,003 | 97,027 | 2,491,030 |

<sup>1</sup> Relative to reference genome
<sup>2</sup> Only found in this individual and not in any of the other 7 genotypes.
\* Reference genome

## Discussion

A major constraint to the use of *M. jansenii* for commercial breeding is the risk of an inedible kernel due to high levels of toxic cyanogenic glycosides. Cyanogenic glycosides have been observed in all the four species of *Macadamia*. However, the concentration varies at different developmental stages (Castada et al., 2020). Even the edible cultivars derived from *M. integrifolia* have genes involved in the cyanogenic glycoside pathway (Nock et al., 2016). However, cyanogenic glycosides levels are extremely low in the kernel of the commercially important species *M. integrifolia* and *M. tetraphylla* (Dahler et al., 1995). The high level of bitterness in the seeds of *M. jansenii* may be associated with high concentrations of cyanogenic glycosides and large numbers of genes for their biosynthesis found in this study. Knowledge of these genes will support efforts to avoid their transfer to domesticated *Macadamia* when using *M. jansenii* as a source of other desirable genes.

Plants may produce antimicrobial proteins as part of their defence against microbial attack. Macadamia seed might have antimicrobial proteins that protect them against attack when germinating in the warm and moist rainforest environment. A new family of antimicrobial peptides, MiAMP-2, was discovered in the seeds of *M. integrifolia* (Marcus et al., 1999). Although only a single gene was found in the *M. jansenii* genome, it encoded a protein with four domains that correspond to the previously reported antimicrobial peptides suggesting that four copies of the peptide could be derived from each translation of this gene. This is the first report of a gene structure for the macadamia anti-microbial peptide with a single intron. This gene has potential for wide use as an antimicrobial protein in plant defence.

Macadamia oil has a unique composition being 75% fat, 80% of which is monounsaturated e.g., oleic oil (C18:1) 55-67%, followed by palmitoleic acid (C16:1) 15-22% (Hu et al., 2019a; Curb et al., 2000; Aquino-Bolaños et al., 2016). The results of analysis of the genes of lipid metabolism in the *M. jansenii* genome are consistent with this fatty acid profile. The number of SAD genes which are responsible for conversion of stearoyl-ACP (18:0) to oleate (18:1) was found to be higher in number than the other genes in these pathways and may explain the desirable high oleic content of macadamias. Retention of these genes will be important in breeding. This species may provide a source of genes for manipulation of lipids in other food crops.

This rare species has a very small population size explaining the low heterozygosity (Ceballos et al., 2018). The heterozygosity was less than one third that of the more widespread, *M. integrifolia*, reported to have a heterozygosity of 0.98% (Topp et al., 2019; Nock et al., 2020). This analysis indicates the importance of conserving the diversity of this endangered species and retaining the unique alleles that may be useful in breeding. *M. jansenii* is a small tree with a high kernel recovery and both of these traits are key for macadamia improvement. Sustainable intensification of production will be facilitated by the breeding of smaller trees and improved kernel recovery is central to kernel yield. Genome level analysis will support field studies for the conservation of this species (Shapcott and Powell, 2011) and molecular analysis of diversity in support of breeding (Mai et al., 2020).

The use of *M. jansenii* as a model in testing genome sequencing and assembly methods (Murigneux et al., 2020; Sharma et al., 2021) is further enhanced by the chromosome level assembly presented here. This is currently the most complete genome sequence available for a macadamia and any member of the more than 1,660 Proteaceae species (Christenhusz and Byng, 2016) making a useful contribution to the goal of sequencing plant biodiversity (Lewin et al., 2018). The Proteaceae belongs to the basal eudicot order Proteales, a sister group to most eudicots (Chanderbali et al., 2016: Drinnan et al., 1994). Among the basal eudicots there are few well characterized genomes. Available genomes include; Aquilegia *coerulea* (Ranuncules) (Filiault et al., 2018), *Papaver somniferum* (Ranuncules) (Pei et al., 2021), Nelumbo nucifera (Proteales) (Ming et al., 2013), *Trochodendron aralioides* (Trochodendrales) (Strijk et al., 2019), *Tetracentron sinense* (Trochodendrales) (Liu et al., 2020). The *M. jansenii* genome provides a valuable contribution to comparative genomics in this group of flowering plants. The chromosome level assembly with an N50 scaffold length of 58 Mb and 96.9% of BUSCO genes compares favourably with those available for other endangered species e.g *Acer yangbiense* with N50 45 Mb and 90.5% BUSCO genes (Giordano et al., 2017), *Ostrya rehderiana* N50 2.31 Mb (Yang et al., 2018) and *Nyssa yunnanensis* with N50 of 985 Kb and BUSCO score of 90.5% (Weixue et al., 2020).

## Experimental procedures

### Plant material

Fresh leaf tissue of *M. jansenii* was collected from *ex-situ* collections of trees at Nambour and Tiaro (three accessions were from the Maroochy Research Facility, Department of Agriculture & Fisheries, Nambour, Queensland, Australia, accessions 1005, 1003 and 1002 and five from Tiaro, Queensland, Australia, Accession #: 1161003, 1161005, 1161001a & 1161001b, 1161004). Fresh leaf tissue (fully expanded young flush) was collected and immediately frozen by placing under dry ice and stored at −80°C until further processed for DNA and RNA extraction.

### DNA and RNA isolation

Leaf tissue was coarsely ground under liquid nitrogen using a mortar and pestle and further ground under cryogenic conditions into a fine powder using a Tissue Lyser (MM400, Retsch, Germany). All accessions were used for DNA isolation. DNA was extracted as per an established method (Furtado, 2014) with minor modification where phenol was excluded from the extraction method. DNA was extracted from 2-3 gm of leaf tissue and dissolved in up to 400 µl of TE buffer.

Accession no. 10051 was used for RNA isolation. RNA was extracted as per established methods (Rubio-Piña and Zapata-Pérez, 2011; Furtado, 2014). RNA was extracted from 2-3 gm of tissue, and treated with extraction buffer, chloroform and phenol/chloropform (1:1) in different steps, followed by further purification using DNase treatment from the Qiagen’s RNeasy Mini kit). RNA quality and quantity were determined using A260/280 and A260/230 absorbance ratio (Nanodrop, Invitrogen USA) and RNA integrity measurements (Bioanalyser, Agilent technology, USA).

### Chromosome level assembly

#### Chicago library sequencing and Sequencing

DNA was isolated as per an established method (Furtado, 2014). Then the library was prepared as described in Putnam et al., (2016). Briefly, ∼500ng of HMW gDNA was reconstituted into chromatin *in vitro* and fixed with formaldehyde. Fixed chromatin was digested with DpnII, the 5’ overhangs filled in with biotinylated nucleotides, and then free blunt ends were ligated. After ligation, crosslinks were reversed, and the DNA was purified from protein. Purified DNA was treated to remove biotin that was not internal to ligated fragments. The DNA was then sheared to ∼350 bp mean fragment size and sequencing libraries were generated using NEBNext Ultra enzymes and Illumina-compatible adapters. Biotin-containing fragments were isolated using streptavidin beads before PCR enrichment of each library. The libraries were sequenced on an Illumina HiSeqX platform to produce 213 million 2×150bp paired end reads, which provided 88.11 x physical coverage of the genome (1-100 kb pairs).

#### Dovetail Hi-C library preparation and sequencing

A Dovetail Hi-C library was prepared in a similar manner as described previously (Lieberman-Aiden et al., 2009). Briefly, for each library, chromatin was fixed in place with formaldehyde in the nucleus and then extracted. Fixed chromatin was digested with DpnII, the 5’ overhangs filled in with biotinylated nucleotides, and then free blunt ends were ligated. After ligation, crosslinks were reversed, and the DNA purified from protein. Purified DNA was treated to remove biotin that was not internal to ligated fragments. The DNA was then sheared to ∼350 bp mean fragment size and sequencing libraries were generated using NEBNext Ultra enzymes and Illumina-compatible adapters. Biotin-containing fragments were isolated using streptavidin beads before PCR enrichment of each library. The libraries were sequenced on an Illumina HiSeqX platform to produce 156 million 2×150bp paired end reads, which provided 3,601.74 x physical coverage of the genome (10-10,000 kb pairs).

#### Scaffolding the assembly with HiRise

The input *de novo* assembly, shotgun reads, Chicago library reads, and Dovetail Hi-C library reads were used as input data for HiRise, a software pipeline designed specifically for using proximity ligation data to scaffold genome assemblies (Putnam et al, 2016). An iterative analysis was conducted. First, Shotgun and Chicago library sequences were aligned to the draft input assembly using a modified SNAP read mapper (http://snap.cs.berkeley.edu). The separations of Chicago read pairs mapped within draft scaffolds were analyzed by HiRise to produce a likelihood model for genomic distance between read pairs, and the model was used to identify and break putative misjoins, to score prospective joins, and make joins above a threshold. After aligning and scaffolding Chicago data, Dovetail HiC library sequences were aligned and scaffolded following the same method. After scaffolding, shotgun sequences were used to close gaps between contigs.

#### Re-sequencing

To study the genetic diversity within the species, re-sequencing of the seven different genotypes was performed on the DNBseq platform (Drmanac et al., 2010). The seven *Macadamia jansenii* samples were selected randomly to represent diversity in the population. A DNBseq library was prepared as follows. Briefly, genomic DNA (1µg) was randomly fragmented using a Covaris, magnetic beads were used to select fragments with an average size of 300-400bp and DNA was quantified using a Qubit fluorometer. The Fragments were subjected to end-repair and 3’ adenylated, adaptors were ligated to the ends of these 3’ adenylated fragments. Then the double stranded products were heat denatured and circularized by the splint oligo sequence, the single strand circle DNA (ssCir DNA) was formatted as the final library. the final library was then amplified to make DNA nanoball (DNB) which had more than 300 copies of each molecule and the DNBs were loaded into the patterned nanoarray. Finally, pair-end 150 bases reads were generated by combinatorial Probe-Anchor Synthesis (cPAS) (MGISEQ-2000).

#### RNA-sequencing

RNA sequencing was undertaken by Macrogen, South Korea. Total RNA was subjected to ribosomal RNA depletion (Ribo zero plant) and then sequenced using Illumina Novaseq 600. Data.

#### Genome assembly quality evaluation & Repetitive element evaluation

The completeness of the genome assembly was evaluated by checking the integrity of the protein coding genes in the Hi-C assembly using Benchmarking Universal Single-Copy Orthologs (BUSCO) (version v5.0.0) analysis with eudicot odb10 dataset with 2326 genes. Repetitive elements in the Hi-C assembly were identified *de novo* and classified using RepeatModeler (version 2.0.1). The repeat library obtained from RepeatModeler was used to identify and mask the repeats in the Hi-C assembly file using RepeatMasker (Version 4.1.0).

#### Structural annotation and functional annotation

The prediction of the protein coding genes in the repeat masked genome was carried out using ab-initio and evidence-based approach. For ab-initio prediction, Dovetail staff used Augustus (version 2.5.5) (Stanke et al., 2006) and SNAP (version 2006-07-28) (Johnson et al., 2008). For evidence based approach, MAKER (Cantarel et al., 2008) was used. For training the ab-initio model for *M. jansenii*, coding sequences from *Malus domestica, Prunus persica* and *Arabidopsis thaliana* were used using AUGUSTUS and SNAP. Six rounds of prediction optimization were done with the package provided by AUGUSTUS. To generate the peptide evidence in Maker pipeline, Swiss-Prot peptide sequences from the UniProt database were downloaded and used in combination with the protein sequences from *Malus domestica, Prunus persica* and *Arabidopsis thaliana*. To assess the quality of the gene prediction AED scores were generated for each of the predicted genes as part of MAKER pipeline. Only those genes which were predicted by both SNAP and AUGUSTUS were retained in the final gene set. To generate the intron hints, a bam file was generated by aligning the RNAseq reads to the genome using the STAR aligner software (version 2.7) and then bam2hints tool was used within the AUGUSTUS. The predicted genes were further characterized for their putative function by performing a BLASTx search against nr protein database (All non-redundant GenBank CDS translations + PDB + SwissProt + PIR+ PRF), as part of annotations undertaken by Dovetail and also by using OmicsBox Ver 1.3.11 (BioBam Bioinformatics, Spain).

#### Gene families

To identify the anti-microbial genes in the genome BLAST homology search was performed to identity transcripts similar to the *M. integrifolia* antimicrobial cDNA (MiAMP2, GenBank: AF161884.1) (Marcus et al., 1999). Then sequence alignment was undertaken using Clone Manager ver 9.0 (SciEd, USA). Multiple Alignment was undertaken using a reference sequence as indicated in the results and alignment parameter scoring matrix of Mismatch (2) Open Gap (4) and Extension-Gap (1).Genes involved in the metabolism of cyanogenic glycosides were identified in the assembly by following a previously described approach (Nock et al., 2016), using BLASTp (1E-5) and sequence homology. Similarly, genes of fatty acid metabolism were identified following the same method.

#### Heterozygosity and genetic diversity analysis

The basic variant analysis (BVA) was performed using Qiagen CLC Genomics Workbench 21.0.4 (CLC bio, Aarhus, Denmark). BGI short read sequences of six genotypes (1003, 1002, 1161003, 1161005, 1161001a, 1161001b) and Illumina reads of one genotype (1005) of *M. jansenii* were mapped to the reference genome of Dovetail Hi-C assembly of *M. jansenii* (1005). Before mapping, the low-quality reads were removed from all the seven genotypes using different CLC trimming parameters (0.05 and 0.01) and the best trimmed reads were selected based upon the Phred score. Then the trimmed reads were mapped against the reference sequence using three different settings: (1.0 LF, 0.95 SF; 1.0 LF, 0.90 SF and 1.0 LF, 0.85 SF), out of which the best mapping was selected and then it was passed through the BVA workflow.

#### Accession numbers

The genome sequence reads, transcriptome sequences and genome assembly of *M. jansenii* have been deposited under NCBI bioproject PRJNA694456.

## Supporting information

fatty

cyano

## Acknowledgements

This project was funded by the Hort Frontiers Advanced Production Systems Fund as part of the Hort Frontiers strategic partnership initiative developed by Hort Innovation, with co-investment from The University of Queensland, and contributions from the Australian Government. We thank the Research Computing Centre (RCC), University of Queensland for support and providing high performance computing resources.

## Author contributions

Contributions of authors were as follows: Designed the study and supervised the project: RJH, AF, BT and MA. Collected sample: MA, BT, AF and PS. Management of germplasm: MA and BT. DNA and RNA isolation: PS and AF. Data analysis and prepared the figures: PS and AF. Bioinformatics analysis: PS, AF, VM, JH and AM. Drafted the manuscript: PS, AF, JH and WT. Data deposition: PS. All authors edited and approved the final manuscript.

## Short legends for Supporting Information

Table S1: Size of each scaffold and number of genes per scaffold

Table S2: SNP heterozygosity statistics in eight *Macadamia jansenii* accessions

Table S3: Genetic diversity statistics in eight *Macadamia jansenii* accessions

Table S4: Genotype-specific polymorphic SNP positions

Table S5: Topologically Associated Domains (TADs) analysis summary

Table S6: TAD statistics at different resolutions

Table S7: Location of fatty acid genes on pseudo-molecules

Table S8: Location of cyanogenic genes on pseudo-molecules

Figure S1: Linkage density histogram of Hi-C assembly of *M. jansenii* genome

Figure S2: BLAST2GO sequence similarity search

Figure S3: Gene ontology (GO) analysis by BLAST2GO

Figure S4: Alignment of the vicilin-like antimicrobial-peptide transcript from *M. integrifolia* and *M. jansenii*

Figure S5: Alignment of anti-microbial CDS sequence of *M. integrifolia* against the *M. jansenii* transcript sequence

Figure S6: Frequency graph of AED scores.

