## Supplementary file for "The genome of the endangered *Macadamia jansenii* displays little diversity but represents an important genetic resource for plant breeding"

**Supplementary tables:**

Table S1: Size of each scaffold and number of genes per scaffold.

| Pseudo-molecule | Size of scaffold | No. of genes per scaffold (Total: 31,591) |
|---|---|---|
| Scaffold # 1 | 67682215 | 2302 |
| Scaffold # 2 | 63669590 | 2089 |
| Scaffold # 3 | 58143993 | 2309 |
| Scaffold # 4 | 56076407 | 2156 |
| Scaffold # 5 | 55220784 | 2154 |
| Scaffold # 6 | 53595462 | 2154 |
| Scaffold # 7 | 52077970 | 2190 |
| Scaffold # 8 | 49563658 | 2291 |
| Scaffold # 9 | 49085581 | 2081 |
| Scaffold # 10 | 48974653 | 2229 |
| Scaffold # 11 | 47698009 | 2120 |
| Scaffold # 12 | 46713600 | 2117 |
| Scaffold # 13 | 45610911 | 2144 |
| Scaffold # 14 | 45288529 | 2227 |
| Total | 739.40 MB | 30563 |
| Total length of assembly | 758.43 Mb | |
| Total genes found | 31,591 | |
| % of genome length in 14 pseudomolecules | 97.5% | |
| % of genes in 14 pseudo-molecules | 96.7% (30563) | |

Table S2: SNP heterozygosity Statistics in eight *Macadamia jansenii* accessions

| Accession ID | Variant alleles: All | Variant alleles: Replacements | Variant alleles: MNV | Variant alleles: Insertions | Variant alleles: Deletions | Variant alleles: SNP | Variant alleles: SNP Homozygous | Variant alleles: SNP Heterozygous | Variant Positions in genome: Homozygous SNPs | Variant Positions in genome: Heterozygous SNPs | Total variant positions in Genome | Variant Positions on genome: Heterozygous SNPs | Heterozygosity |
|---|---|---|---|---|---|---|---|---|---|---|---|---|---|
| 1005* | 5,418,086 | 7,642 | 158,763 | 287,946 | 198,900 | 4,764,835 | 4,019 | 4,735,918 | 4,019 | 2,428,956 | 2,432,975 | 2,428,956 | 0.31 |
| 1161004 | 5,415,612 | 6,534 | 129,887 | 218,716 | 158,864 | 4,901,611 | 784,323 | 4,013,541 | 784,323 | 2,038,553 | 2,822,876 | 2,038,553 | 0.26 |
| 1161003 | 6,785,189 | 10,754 | 184,115 | 320,586 | 235,055 | 6,034,679 | 1,047,938 | 4,851,037 | 1,047,938 | 2,465,089 | 3,513,027 | 2,465,089 | 0.32 |
| 1161005 | 6,204,994 | 9,908 | 175,182 | 307,401 | 224,149 | 5,488,354 | 780,593 | 4,612,495 | 780,593 | 2,347,362 | 3,127,955 | 2,347,362 | 0.30 |
| 1161001a | 6,977,842 | 11,107 | 195,865 | 335,236 | 239,389 | 6,196,254 | 875,565 | 5,211,753 | 875,565 | 2,649,035 | 3,524,600 | 2,649,035 | 0.34 |
| 1003 | 7,050,861 | 11,344 | 199,289 | 341,519 | 244,482 | 6,254,227 | 891,669 | 5,256,600 | 891,669 | 2,672,103 | 3,563,772 | 2,672,103 | 0.34 |
| 1002 | 6,759,260 | 11,299 | 188,202 | 339,398 | 246,936 | 5,973,425 | 1,044,208 | 4,812,812 | 1,044,208 | 2,447,418 | 3,491,626 | 2,447,418 | 0.31 |
| 1161001b | 6,704,384 | 10,277 | 183,381 | 319,372 | 228,920 | 5,962,434 | 824,632 | 5,027,801 | 824,632 | 2,556,695 | 3,381,327 | 2,556,695 | 0.33 |
| | | | | | | | | | 6,252,947 | 19,605,211 | 25,858,158 | 2,450,651 | 0.31 |

*Reference genome

Table S3: Genetic diversity statistics in eight *Macadamia jansenii* accessions

| | Number of variants that are heterozygous and homozygous | Number of variants that are heterozygous only | Number of variants that are heterozygous or homozygous | Number of variants that are homozygous only | Total positions that are polymorphic | Genome size | Genetic Diversity (%) |
|---|---|---|---|---|---|---|---|
| All genotypes | 6,856,125 | 5,253,468 | 1,602,657 | 350,149 | 7,206,274 | 780,000,000 | 0.92 |
| Accession ID | | | | | | | |
| 01 1005* | 2,249,732 | 1,496,986 | 752,746 | 3,070 | 2,252,802 | 780,000,000 | 0.29 |
| 02 1161004 | 1,902,705 | 1,082,034 | 820,671 | 111,100 | 2,013,805 | 780,000,000 | 0.26 |
| 03 1161003 | 2,306,771 | 1,184,768 | 1,122,003 | 162,541 | 2,469,312 | 780,000,000 | 0.32 |
| 04 1161005 | 2,190,169 | 1,160,412 | 1,029,757 | 85,258 | 2,275,427 | 780,000,000 | 0.29 |
| 05 1161001a | 2,484,209 | 1,381,312 | 1,102,897 | 109,875 | 2,594,084 | 780,000,000 | 0.33 |
| 06 1003 | 2,505,726 | 903,069 | 1,602,657 | 113,837 | 2,619,563 | 780,000,000 | 0.34 |
| 07 1002 | 2,286,675 | 1,153,549 | 1,133,126 | 165,048 | 2,451,723 | 780,000,000 | 0.31 |
| 08 1161001b | 2,394,003 | 1,347,285 | 1,046,718 | 97,027 | 2,491,030 | 780,000,000 | 0.32 |

*Reference genome

Table S4: Genotype-specific polymorphic SNP positions

| | Heterozygous position in only one genotype | Homozygous position in only one genotype | Total number of Heterozygous and homozygous positions in only one genotype |
|---|---|---|---|
| All genotypes | 2,400,562 | 144,304 | 2,544,866 |
| Accession ID | | | |
| 01 1005* | 585,053 | 762 | 585,815 |
| 02 1161004 | 187,441 | 8,615 | 196,056 |
| 03 1161003 | 521,184 | 44,014 | 565,198 |
| 04 1161005 | 608,485 | 40,629 | 649,114 |
| 05 1161001a | 95,089 | 8,155 | 103,244 |
| 06 1003 | 99,715 | 9,878 | 109,593 |
| 07 1002 | 219,933 | 27,408 | 247,341 |
| 08 1161001b | 83,662 | 4,843 | 88,505 |

*Reference genome

Table S5: Topologically Associated Domains (TADs) analysis summary

| Feature | Count |
|---|---|
| A Compartments | 37 (49.77%) |
| B Compartments | 38 (49.36%) |
| TADs at 10 kbp resolution | 13 |
| TADs at 25 kbp resolution | 84 |
| TADs at 50 kbp resolution | 76 |
| Isochores | 6,473 |
| CTCF Binding Sites | 2,423 |

Table S6: TAD statistics at different resolutions

| Resolution (kbp) | Number of TADs | Mean TAD Size (bp) | Basepairs in TADs (kbp) | Percent genome contained in TADs |
|---:|---:|---:|---:|---:|
| 10 | 13 | 348,461 | 45 | 0.61% |
| 25 | 84 | 497,619 | 418 | 5.65% |
| 50 | 76 | 1,553,947 | 1,123 | 15.19% |

Table S7: Location of fatty acid genes on pseudo-molecules (Supplied as separate excel sheet).

Table S8: Location of cyanogenic genes on pseudo-molecules (Supplied as separate excel sheet).

**Supplementary figures:**

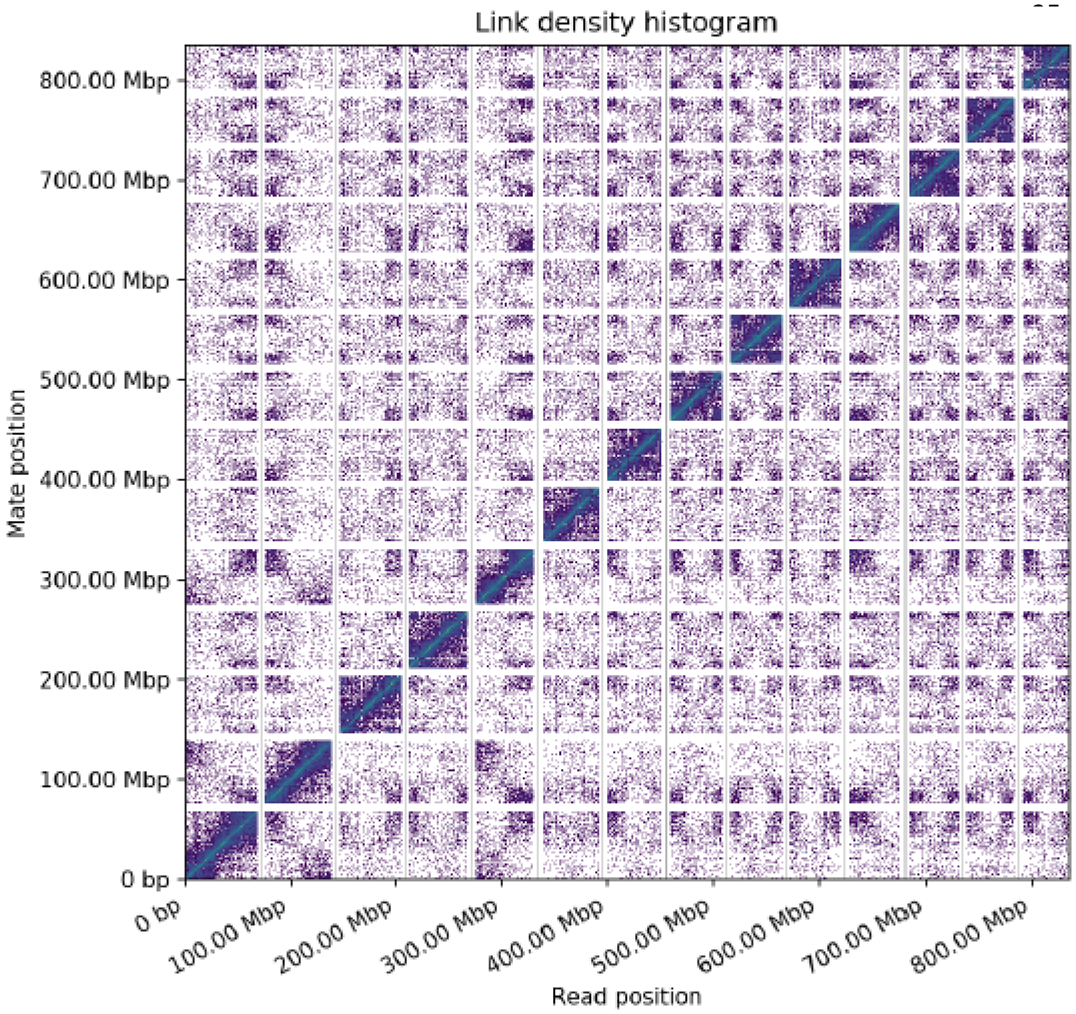


Figure S1: Linkage density histogram of Hi-C assembly of *M. jansenii* genome

The x and y axes give the mapping positions of the first and second read in the read pair respectively, grouped into bins. The colour of each square gives the number of read pairs within that bin. White vertical and black horizontal lines have been added to show the borders between scaffolds. Scaffolds less than 1 Mb are excluded.

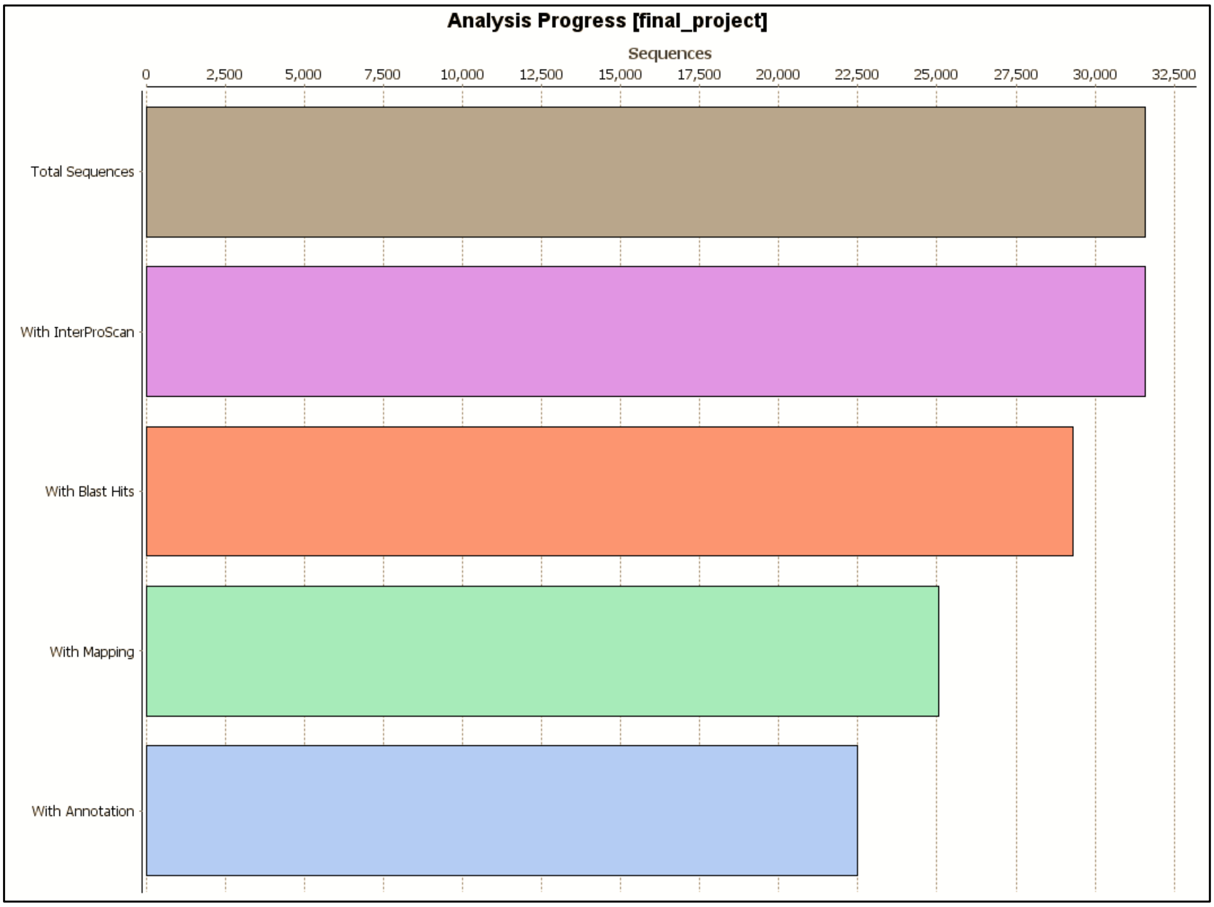


Figure S2**:** BLAST2GO sequence similarity search

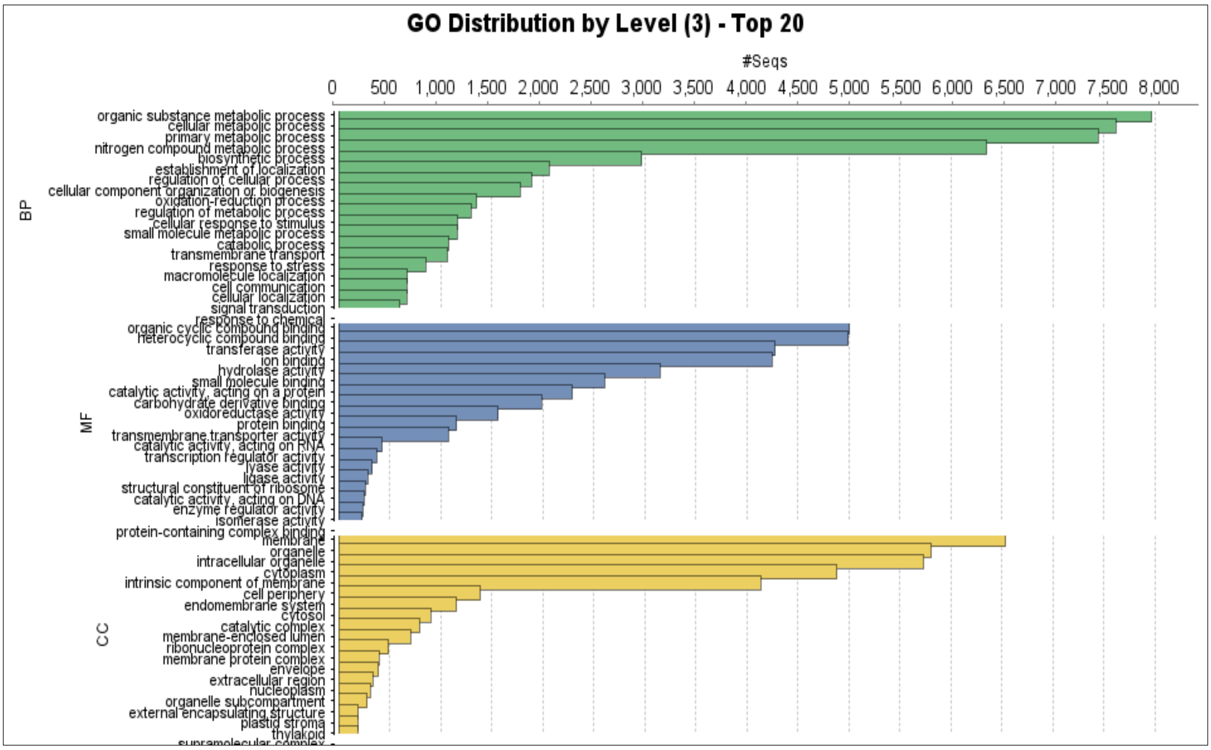


Figure S3: Gene ontology (GO) analysis by BLAST2GO

| **A** Sequence Details | Start | End | Global alignment of transcript compared to MiAMP-2 as reference | | | % of transcript matched |
|---|---|---|---|---|---|---|
| | | | bases matched | bases not matched | % match | |
| **MiAMP-2** | 1 | 2182 | | | | |
| **ANN01396** | 1 | 1998 | 1962 | 220 | 89.92 | 98.20 |
| **ANN01394** | 1 | 960 | 600 | 1605 | 27.50 | 62.50 |
| **ANN01395** | 1 | 348 | 262 | 1921 | 12.01 | 75.29 |
| **ANN02865** | 1 | 309 | 233 | 1950 | 10.68 | 75.40 |
| **ANN08351** | 1 | 1062 | 678 | 1520 | 31.07 | 63.84 |
| **ANN09965** | 1 | 1938 | 991 | 1329 | 45.42 | 51.14 |
| **ANN15809** | 1 | 1863 | 980 | 1368 | 44.91 | 52.60 |
| **ANN23835** | 1 | 1590 | 900 | 1326 | 41.25 | 56.60 |

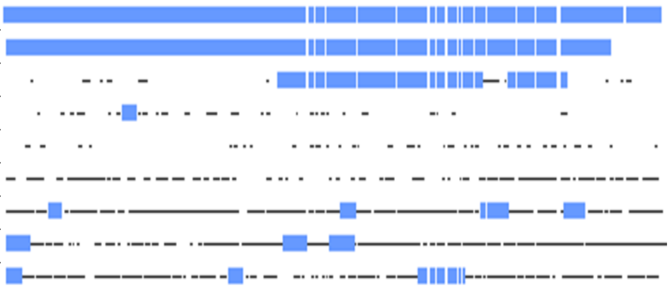

| **B** Sequence Details | Start | End | Global alignment of transcript compared to ANN01396 as reference | | | % of transcript matched |
|---|---|---|---|---|---|---|
| | | | bases matched | bases not matched | % match | |
| **ANN01396** | 1 | 1998 | | | | |
| **ANN01394** | 1 | 960 | 594 | 1427 | 29.73 | 61.88 |
| **ANN01395** | 1 | 348 | 261 | 1738 | 13.06 | 75.00 |
| **ANN02865** | 1 | 309 | 234 | 1764 | 11.71 | 75.73 |
| **ANN08351** | 1 | 1062 | 672 | 1340 | 33.63 | 63.28 |
| **ANN09965** | 1 | 1938 | 971 | 1183 | 48.60 | 50.10 |
| **ANN15809** | 1 | 1863 | 960 | 1231 | 48.05 | 51.53 |
| **ANN23835** | 1 | 1590 | 874 | 1214 | 43.74 | 54.97 |

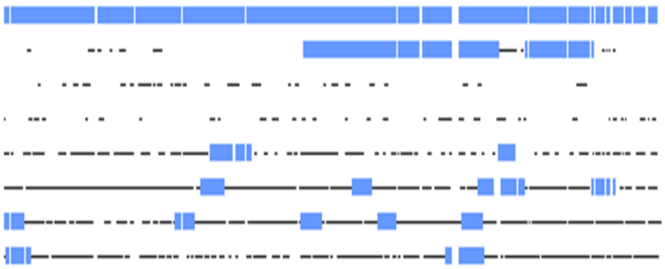

Figure S4: Alignment of the vicilin-like antimicrobial-peptide transcript from *M. integrifolia* and *M. jansenii*

**Figure S4 (A):** MiAMP-2, antimicrobial cDNA sequence from *Macadamia integrifolia*; ANN01396, ANN01395, ANN01396, ANN08351, ANN02865, ANN23835, ANN15809, ANN09965, transcripts identified from the functional annotation of the *Macadamia jansenii* assembly. The MiAMP-2 sequence show high homology to two of the *M. jansenii* transcripts, ANN01396 and ANN01394. **Figure S4 (B):** The *M. jansenii* transcript ANN01396 shown partial homology to the ANN01394 transcript mainly at the second half of the sequence.

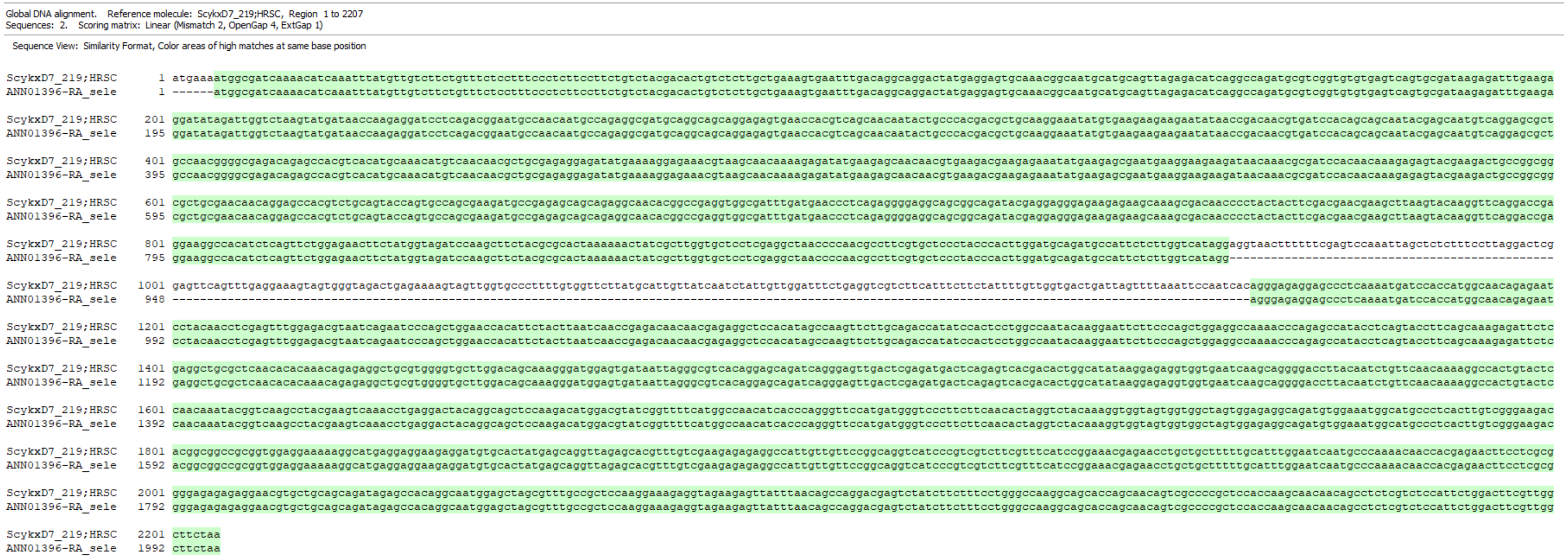


Figure S5: Alignment of anti-microbial CDS sequence of *M. integrifolia* against the *M. jansenii* transcript sequence. The gap in between depicts the intron position in the gene.

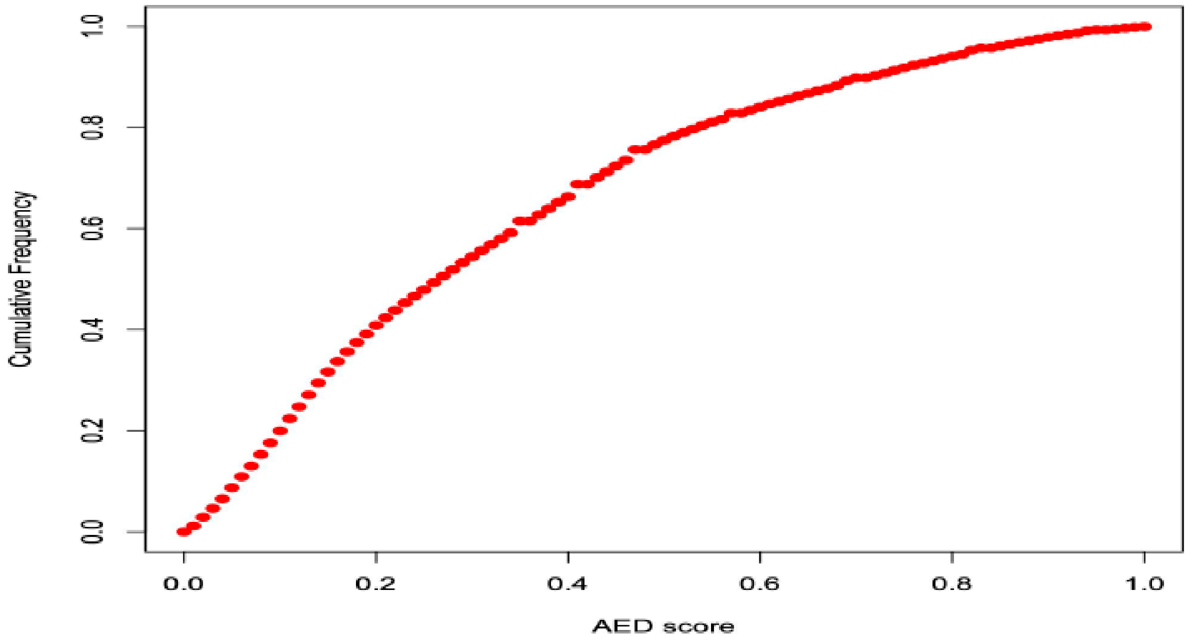


Figure S5: Frequency graph of AED scores.

Annotation edit distance (AED) is a general measure of how well the predicted gene is supported by external evidence (UniProt protein and mRNA sequences). AED score ranges from 0 to 1 and a lower score represents more evidence support for the gene. AED is calculated for every gene. The AED cumulative frequency graph above provides an overview of the quality of the gene annotation.
